# Older Adults Forage Broadly for Information Despite Cost

**DOI:** 10.64898/2026.09.14.751536

**Authors:** Abigail Hedden, David L. Barack, Akram Bakkour

**Affiliations:** Department of Psychology, University of Chicago, Chicago, IL; Institute for Mind and Biology, University of Chicago, Chicago, IL; Neuroscience Institute, University of Chicago, Chicago, IL; Department of Philosophy and Department of Social and Decision Sciences, Carnegie Mellon University, Pittsburgh, PA; Center for Philosophy of Science, University of Pittsburgh, Pittsburgh, PA

**Keywords:** Aging, Learning, information foraging, search

## Abstract

Healthy aging has long been associated with impairments in learning from rewards. However, typical paradigms do not adequately distinguish between being motivated by information to learn about the world and being motivated by rewarding outcomes from our decisions. The present study describes how these distinct motivations guide learning about the environment across the lifespan. Participants (N=169; ages 18-71 [M=39, SD=15.4]) completed a task based on the board game Battleship, making sequential choices about which tile to uncover on a grid with the goal of finding and learning hidden shapes. While older adults performed worse on the task in terms of overall score, their search behavior stabilized earlier. Older adults also preferred to seek information over reward, spreading their choices more widely, making more choices per trial, and acquiring more information per trial. Additionally, older adults showed a persistent reliance on information gained from the outcomes of their choices. These findings suggest that, as individuals age, they prefer to seek information, which earns them fewer points and slows learning but incurs no cost to long-run projected performance. Despite the conventional view linking age with diminished learning, in a complex environment that disentangled information and reward, older adults remained engaged with no projected learning deficits. Our findings challenge assumptions about cognitive decline in aging and reframe information seeking in older adults as a persistent motivational preference that need not serve learning, raising the question of why the drive to seek information intensifies with age even when it confers no advantage.

## Introduction

Older adults are notoriously impaired when learning from rewards (Lerner et al., 2018; Marschner et al., 2005; Mell et al., 2005). However, many different motivations drive learning in humans, including seeking information about the environment. Unlike rewards, the search for and use of information across the lifespan remains little investigated. Using a structure learning task that disambiguates reward-from information-seeking, we discovered that older adults show a pronounced, persistent preference for information over reward. While older adults’ preference for information and increased directed exploration came at a cost to earnings and learning speed, there were no projected longer-term performance deficits and no assessed deficits of age itself.

Both reward and information can motivate learning about the environment. By reward, we mean points, juice, food, or other valuable outcomes from choices, and by information, we mean the outcomes from choices that are predictive of receiving valuable outcomes in the future. Both reward and information are used to estimate the value of options. To illustrate, suppose you are new in town and must decide on a restaurant for dinner. You could visit some restaurant and try items on the menu; the food provides rewards to use to update your value of the restaurant. Alternatively, you could seek information by reading reviews on the internet, which would also allow you to update your estimate of the restaurant’s value. Here, we ask whether older adults are impaired at learning a new environment when given the opportunity to gather information in addition to experiencing reward.

A range of changes in reward-related motivation may explain previously observed deficits in learning in older adults. These include challenges in using optimal strategies (Mata et al., 2011) or in adapting to changing reward-contingencies in their environment (Bartus et al., 1979; Mell et al., 2005). Such impairments may also arise from reduced activity in the neural circuit responsible for reward predictions (Samanez-Larkin et al., 2014), shifts in reward sensitivity (Frank & Kong, 2008), or changes in preferences (Eppinger et al., 2012). Despite variability in tasks, methods, and findings, the consensus is that older adults’ abilities to learn about novel environments from feedback is impaired. The findings from the current study challenge this consensus.

While past studies largely focused on the impact of aging on learning from rewards, humans and non-human animals also seek information to learn about the environment (Barack et al., 2023). When learning novel environments, people may seek information with instrumental utility that can be used to gather reward (Dezza et al., 2022) or non-instrumental information that cannot be used to gather rewards in the future (Bromberg-Martin & Monosov, 2020). Seeking non-instrumental information may reflect hard-wired preferences for gathering information derived from instrumental contexts in the selective history of organisms. A rich literature has explored the value of non-instrumental information in decision making in a range of species (typically in the form of so-called ‘observing responses’, Blanchard et al., 2015; Bromberg-Martin & Hikosaka, 2009, 2011; Bromberg-Martin & Monosov, 2020; Charpentier et al., 2018; Wyckoff, 1952). Research has also focused on how humans seek instrumental information, information that can be put to adaptive use (Dezza et al., 2022; Howard, 1966; Kobayashi & Hsu, 2019; Wilson et al., 2014). This includes computational and behavioral investigations of sense-making (Wojtowicz et al., 2022; Wojtowicz & DeDeo, 2020), curiosity (Kidd & Hayden, 2015; Murayama, 2022; Murayama et al., 2019; Oudeyer, 2018; Sharot & Sunstein, 2020) or interest (Berlyne, 1949; Peterson & Hidi, 2019), intrinsic motivation (Baranès & Oudeyer, 2009; Bennett et al., 2016; Gottlieb et al., 2013, 2016; Gottlieb & Oudeyer, 2018; Oudeyer & Baranes, 2008), and the drive to reduce uncertainty (Markant et al., 2013; Markant & Gureckis, 2010, 2012; Meder et al., 2022; Nelson, 2005; Nelson et al., 2018; Oaksford & Chater, 1996), as well as neuroscientific investigations of the mechanisms underlying the search for such information (Kobayashi & Kable, 2024; Li et al., 2022).

How does the search for useful information change across the lifespan? Older adults are slower in their search for information (Sharit et al., 2015) and demonstrate poorer performance than younger adults on a simulated customer service task (Czaja et al., 2001) and a stock picking task (Mata et al., 2010). However, once they have learned a task, older adults perform comparably to younger people despite slower response times (Sharit et al., 2015), suggesting any performance differences become nonsignificant with sufficient practice (Czaja et al., 2001). Task structure also influences information search: the reliance on prior knowledge (Blanco et al., 2016) can harm older adults’ learning, but this is possibly offset by the helpful utilization of simpler strategies (Mata et al., 2007, 2010). While the ability to search for and learn from information in older adults has been previously studied, this past work failed to properly disambiguate information from reward and often concerns explicit searches for information before making decisions (e.g., Mata et al., 2010). Sequential decisions to search for information followed by seeking reward may be useful in the laboratory. In contrast, decisions in the real world often simultaneously involve both reward and information. How does aging impact the search for reward and information during decision-making?

We examined how healthy adults across the lifespan utilize information and reward to learn about the environment using a shape learning task. We predicted that younger adults would use reward and information outcomes differently during learning, prioritizing reward over information during learning. In contrast, we predicted that although older adults would learn the environment, their impaired ability to learn from rewards would lead to a greater reliance on information. We found that older adults search far more actively for information but earn fewer points overall and learn shapes more slowly when compared to younger adults but are projected to reach the same level of performance in the long-term. Older adults prioritize information over reward during search, even in later trials when no longer useful. Younger adults, in contrast, switch from prioritizing information early in the experimental session to prioritizing reward later in the same session. Our findings reveal a pronounced, persistent drive to seek information in aging that is detrimental to earning reward and to the speed of shape learning, raising the question of why older adults invest so heavily in information search that does not pay off. The discovery of enhanced, non-instrumental information seeking in older adults promises a better understanding of real-world behavior in aging populations.

## Results

To understand how humans use information and reward to learn the structure of environments, we employed a shape search task (Barack et al., 2023). In this task (**Figure 1**), participants searched for and learned five shapes across many decisions and trials. As opposed to traditional feedback-based reinforcement learning tasks, participants were able to use two distinct motivations, information and reward, to learn the shapes and their location in this novel environment.

**Figure 1.**
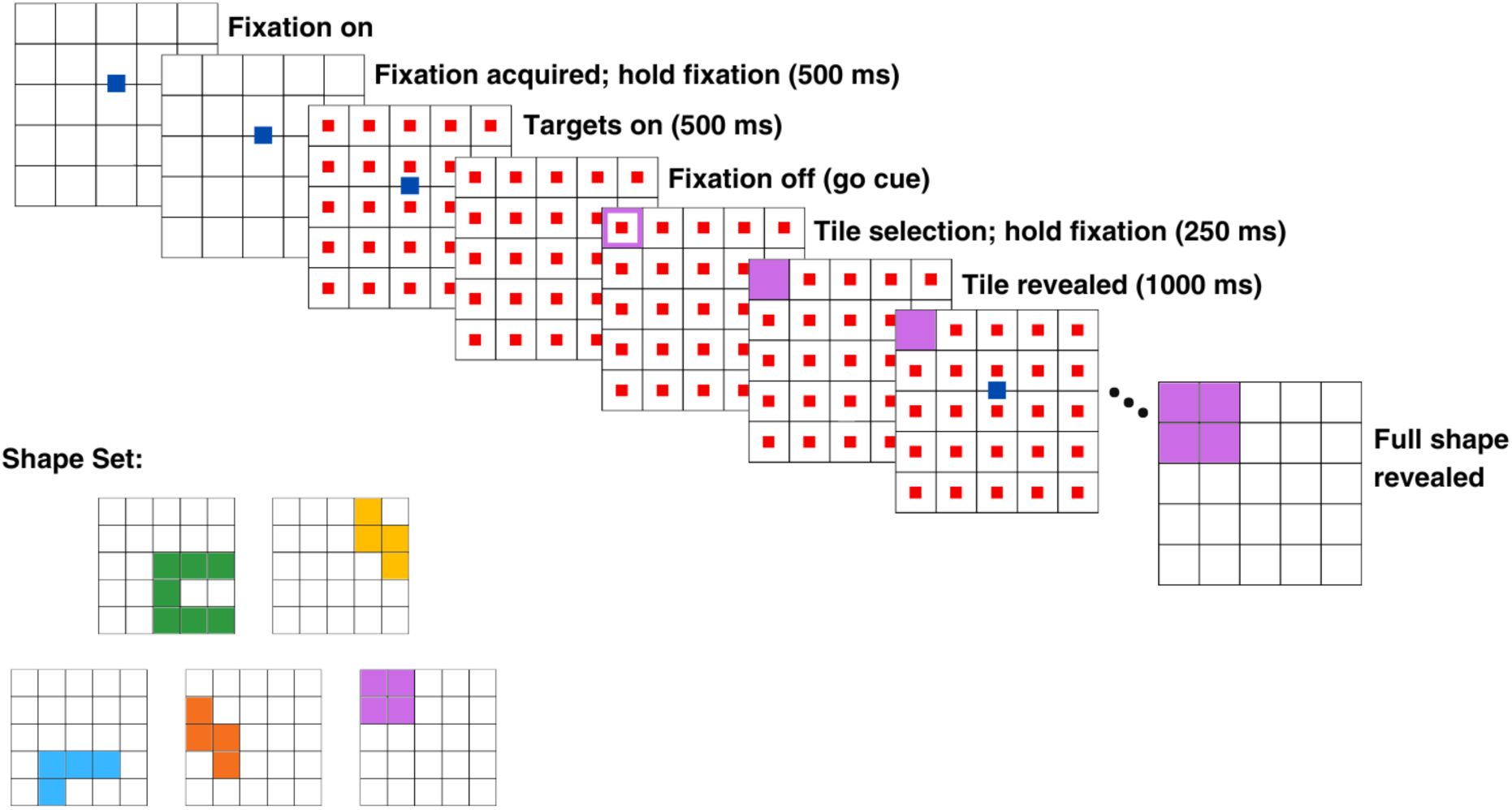
Battleship Task. In each trial, one shape from the 5-shape set (bottom left) was hidden on a 5×5 grid of smaller squares (tiles). For each choice, participants placed the mouse cursor over the central fixation (blue square) for 500 ms in order to reveal the possible tile choices (red squares). Placing the cursor over a tile for 250ms ‘flips’ the tile over to reveal what is under it (a white tile [not part of a shape] or a colored tile [part of a shape]). After revealing all tiles that made up a shape, the entire grid revealed itself, showing the full shape, before a new trial started.

*Older adults perform worse, gather more information, and stabilize behavior sooner* Older adults are impaired at learning from reward feedback (Eppinger et al., 2013; Frank & Kong, 2008; Lerner et al., 2018; Mell et al., 2005). Consistent with these impairments, we find that, as our participants’ age increases, they earn fewer points overall by the end of the fixed 50-minute experimental session (ordinary least squares (OLS) regression β =-0.961, 95% confidence interval (CI) [-1.723,-0.200], *p* = 0.014).

We next examined how older and younger adults searched for shapes. A linear regression of age against the total number of completed trials revealed that the number of trials completed in the task decreased as age increased (OLS β =-0.200, 95% CI [-0.365,-0.035], *p* = 0.018). This was not simply due to older adults performing the task more slowly, as the speed at which the mouse cursor moved across the screen did not differ with age (mixed-effects linear regression β =-2.338 x 10^-4^, 95% CI [-5.408 x 10^-4^, 7.316 x 10^-5^], *p* = 0.135). There was also no relationship between the total number of choices made in the task and age — older adults did not make more choices overall than their younger counterparts (OLS β = 0.278, 95% CI [-0.563, 1.119], *p* = 0.516). There was, however, a positive relationship between the average number of choices per trial and age, where older age was associated with making more choices per trial (OLS β = 0.030, 95% CI [0.017, 0.044], *p* < 0.001).

The benefit to making more choices per trial in older age becomes evident when considering the amount of information gained from those choices. Information in this case is defined as the amount of uncertainty about the hidden shape that is reduced by making a choice (see Methods). A linear regression revealed that, as age increased, the average amount of information per trial increased (β = 0.023, 95% CI [0.012, 0.033], *p* < 0.001). Although older adults completed fewer trials, resulting in fewer points overall, they tended to make more choices per trial and gathered more information from those choices.

The task structure also allowed us to investigate when participants stabilized their search behavior. Here, stabilization of search behavior does not identify when participants stop learning, but rather when the rate at which their behavior changes has become stable. To determine when each participant’s search behavior stabilized, we performed a changepoint detection test on the mean and variance of choices. As participants learn the shape set, both the mean number of choices to find the shape and the variance in the number of choices should decrease (see Methods). The last changepoint (either by variance or by mean) for each participant marks the point at which their search behavior stabilized. The changepoint detection test failed to detect a changepoint for 13 of 169 (∼8%) participants. A logistic model indicated no significant relationship between age and the likelihood of detecting a changepoint (odds ratio (OR) = 1.025, 95% CI [0.99, 1.07], *p*= 0.199), and these participants were excluded from the following analyses. We found that the trial number on which the final changepoint was detected decreased with age (OLS β =-0.267, 95% CI [-0.471,-0.064], *p* = 0.010). In other words, older adults’ search behavior stabilized earlier than younger adults’.

### Older adults sample the board more broadly with more targeted exploration

Older adults made more choices per trial, gathered more information on each trial, completed fewer trials overall, stabilized their search behavior sooner, and attained lower total scores compared to young adults. To better understand this pattern of results, we next investigated how adults sampled the board, focusing on the spread of their choices as a proxy of their exploration of the novel environment. We employed mixed-effects linear regression to regress Euclidean pairwise choice distance (see Methods) against age, choice number in the task, trial number in session, and all interactions (**Figure 2**). All effects are outlined in **Table 1**.

**Figure 2.**
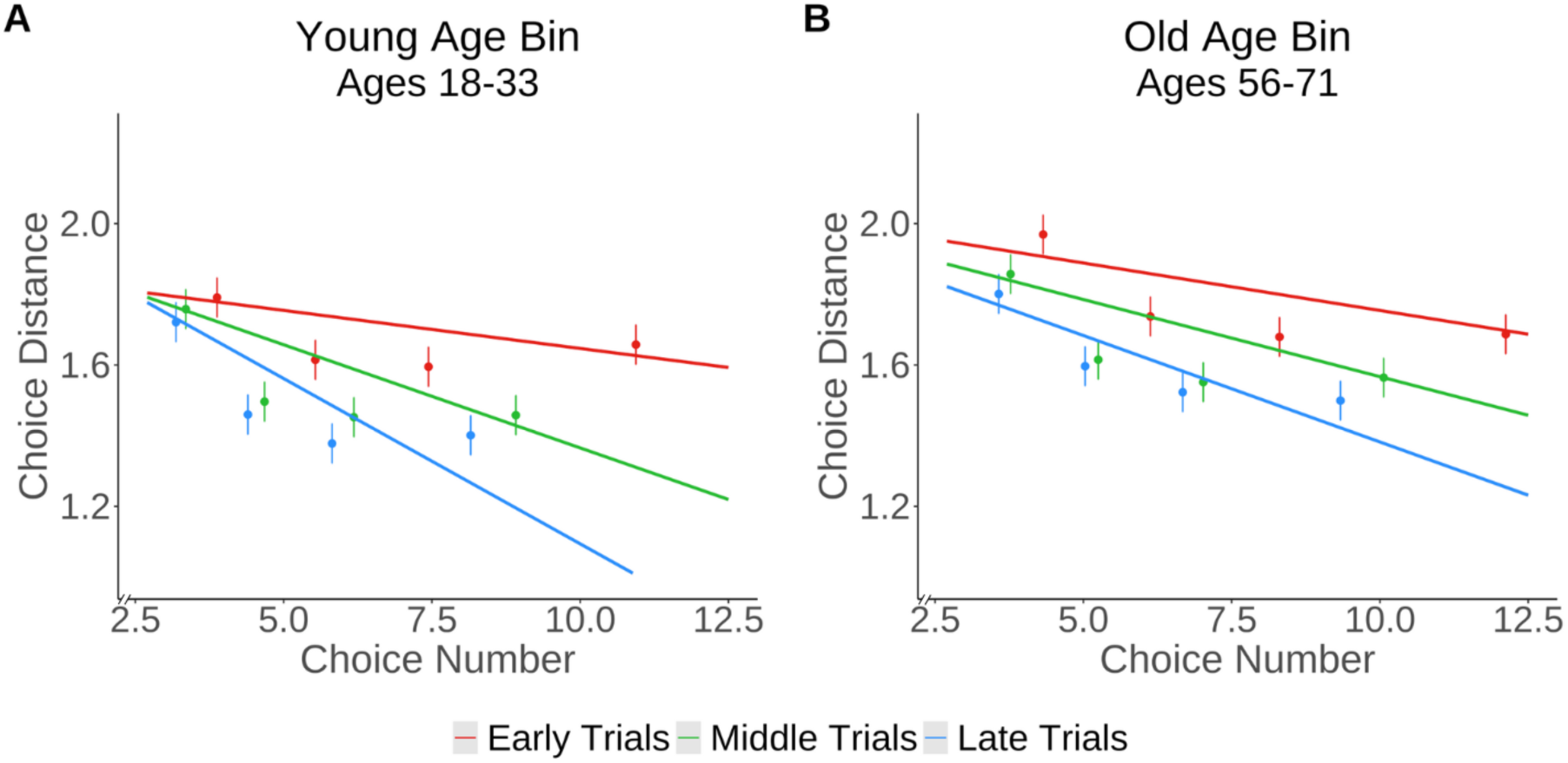
Younger Adults Narrow Their Choices Over Time, While Older Adults Maintain a Broader Search. Choice distance — the mean pairwise Euclidean distance between successively chosen tiles — is plotted against choice number within a trial in **A)** the youngest (ages 18-33), and **B)** the oldest (ages 56-71) participants. Trial data were binned into 3 bins, resulting in early, middle, and late trials. Within each trial bin, choice numbers were binned into 5 equal-sized bins. Dots represent the mean choice distance across participants within each choice number bin within each trial bin. Lines represent the marginal effects of choice number on choice distance for each age bin. Red lines/dots indicate early trials, green lines/dots indicate middle trials, and blue lines/dots indicate late trials.

**Table 1.** Choice Distance Mixed-Effects Model Results. The linear effects of age, trial number, choice number, and all interactions on choice distance were examined.

| *Predictors* | **Choice Distance** |  |  |
| --- | --- | --- | --- |
|  | *Estimates* | *CI* | *p* |
| (Intercept) | 1.68 | 1.59 – 1.77 | **<0.001** |
| Age | 0.01 | 0.00 – 0.01 | **<0.001** |
| Choice Number | 0.01 | -0.00 – 0.01 | 0.191 |
| Trial | 0.01 | 0.01 – 0.01 | **<0.001** |
| Age × Choice Number | $-3.80 \times 10^{-4}$ | -0.00 – -0.00 | **<0.001** |
| Age × Trial | $-1.21 \times 10^{-4}$ | -0.00 – -0.00 | **<0.001** |
| Choice Number × Trial | $-2.00 \times 10^{-3}$ | -0.00 – -0.00 | **<0.001** |
| Age × Choice Number × Trial | $1.83 \times 10^{-5}$ | 0.00 – 0.00 | **<0.001** |
| **Random Effects** |  |  |  |
| $\sigma^2$ | 0.69 | | |
| T<sub>00</sub> Subject | 0.03 |  |  |
| ICC | 0.04 |  |  |
| N<sub>Subject</sub> | 169 |  |  |
| Observations | 77666 |  |  |
| Marginal R<sup>2</sup>/Conditional R<sup>2</sup> | 0.036 / 0.074 |  |  |

We discovered a main effect of age on distance. As age increased, the distance between successive chosen tiles also increased (see “Age” in **Table 1**), indicating that older adults spread their choices more broadly across the board (i.e., their choices were less clustered) than younger adults. Significant interactions between both age and choice number (see “Age ✕ Choice Number” in **Table 1**) and age and trial number (see “Age ✕ Trial Number” in **Table 1**) revealed that the impact of increasing choice number or increasing trial number on choice distance diminished as age increased.

This analysis also revealed a significant interaction between choice number and trial number (see “Choice Number ✕ Trial Number” in **Table 1**). Initially, in early trials, participants made widely spaced choices, resulting in a broad sampling of the board. As the trial progressed (i.e., as choice number increased) and participants uncovered portions of the shape, their choices became more clustered, resulting in smaller distances between selections. In later trials, after participants had learned the shapes and their locations, the distance between their choices decreased much more rapidly as choice number increased. This effect, however, varied with age. A three-way interaction between age, choice number, and trial number (see “Age ✕ Choice Number ✕ Trial Number” in **Table 1**) revealed that the effect of choice number and trial number on choice distance was weaker in older adults. In other words, the rapid decrease in choice distance as choice number progressed in later trials was less pronounced in older adults compared to younger adults. Older adults exhibited a more consistent spread in their choices throughout the trials, regardless of choice number. In contrast, younger adults showed a more rapid decrease in the distance between their choices as trials progressed.

Older adults spread their choices more broadly across the board, even later in sessions, indicating consistently greater exploration in this task. However, was this exploration random, or did it reflect a directed strategy? We reasoned that movement consistent with the expected distance (see Methods) indicated random exploration, whereas greater deviations from the expected distance reflected directed exploration. Deviation from expectation was therefore used as a proxy for directed exploration. To investigate random vs. directed exploration, we ran the same mixed-effects regressions for these measures as we did for choice distance. There was no three-way interaction against random exploration (Age ✕ Choice Number ✕ Trial β = 0.0022, 95% CI [−0.001, 0.006], p = 0.21), but a significant three-way interaction against directed exploration (Age ✕ Choice Number ✕ Trial β = 0.0088, 95% CI [0.006, 0.012], p < 0.0001). Age was associated with preserved spreading of choices throughout a trial even late in the game, indicating persistent directed exploration in older adults.

### Information is prioritized over reward in older adults

We found that, when searching for shapes, older adults persistently spread their choices more widely across the board and exhibit more directed exploratory behavior. How might the choice of tiles be related to reward and information? To address this question, we explored the impact of expected reward and expected information on choice. Expected reward represented the frequency with which a particular tile had yielded a reward relative to the total number of times it had been selected (see Methods). Expected information for a given tile was defined as the mean in the change in entropy computed over a Dirichlet distribution if the next tile were a hit or a miss (see Methods). Different tiles were associated with different probabilities of leading to reward based on prior choice outcomes and different levels of information.

We performed a mixed-effects linear regression to regress choice number in trial against trial number in session, the expected reward for the chosen tile, the expected information for the chosen tile, age, and all interactions (**Figure 3**). There were main effects for information, reward, and trial number on choice number (**Table 2**). As choice number increased, the impact of expected information on tile selection decreased (see “Information” in **Table 2**), indicating that participants were choosing more informative tiles earlier in trials. Moreover, the impact of reward on tile selection increased with increasing choice number (see “Reward” in **Table 2**), indicating that participants chose more rewarding tiles later in trials. These effects were made clearer upon examining interactions between these variables. A significant interaction between information and reward was found (see “Information ✕ Reward” in **Table 2**). This interaction is unsurprising, given the negative relationship between information and reward across choices (β =-0.021, 95% CI [-0.032,-0.009], p < 0.001). A significant interaction was also found between trial number and age (see “Trial ✕ Age” in **Table 2**). This suggests that the number of choices per trial decreased more quickly as trials progressed in younger adults when compared to older adults. A three-way interaction was also found between information, reward, and trial number (see “(Information ✕ Reward) ✕ Trial” in **Table 2**). This effect was expected, as information and reward became more negatively correlated as trial number increased (β =-3.069 x 10^-3^, 95% CI [-3.762 x 10^-3^,-2.376 x 10^-3^], p < 0.0001), a pattern consistent with the participants’ learning of the shapes across trials. At the end of the task, the first choice in a trial is maximally informative, as it likely reveals which of the learned shapes is hidden. The subsequent choices within the trial thus become less informative and more rewarding, leading to the observation of an increasingly negative relationship as the trial number progresses.

**Figure 3.**
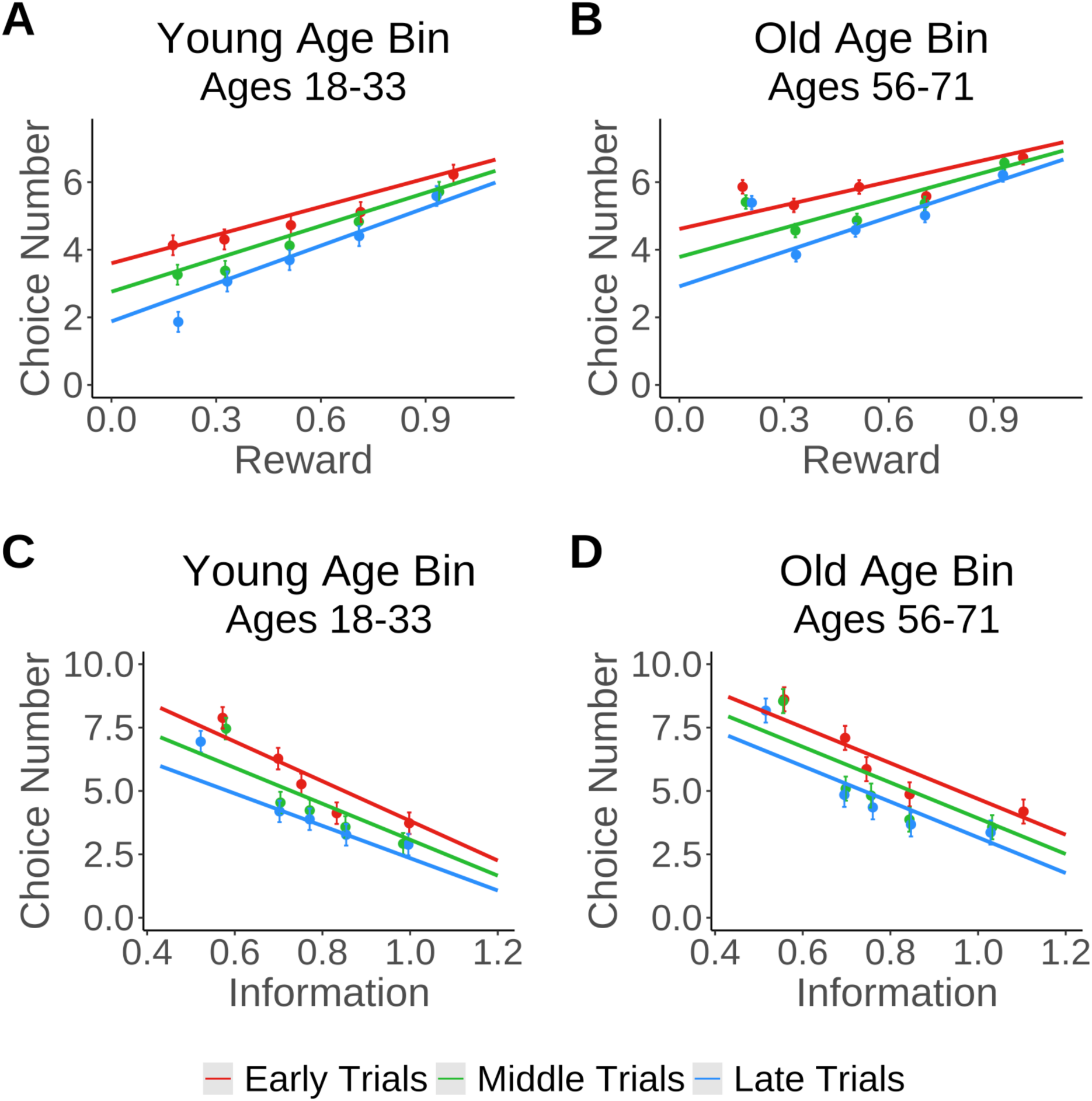
Older Adults Maintain Prioritization of Information, While Younger Adults Decrease Prioritization of Information as They Learn the Shapes. Choice number within a trial is plotted against Reward **(A-B)** and Information **(C-D)** in the youngest **(A,C)** and oldest **(B,D)** adults in our sample. Trial numbers were binned into 3 bins, resulting in early, middle, and late trials. Within each trial bin, reward/information data were binned into 5 equal-sized bins. In each plot, dots represent the mean choice numbers across participants within each reward/information bin within each trial bin. Lines represent the marginal effects of information/reward on choice number for each trial bin. Red lines/dots indicate early trials, green lines/dots indicate middle trials, and blue lines/dots indicate late trials.

**Table 2.**
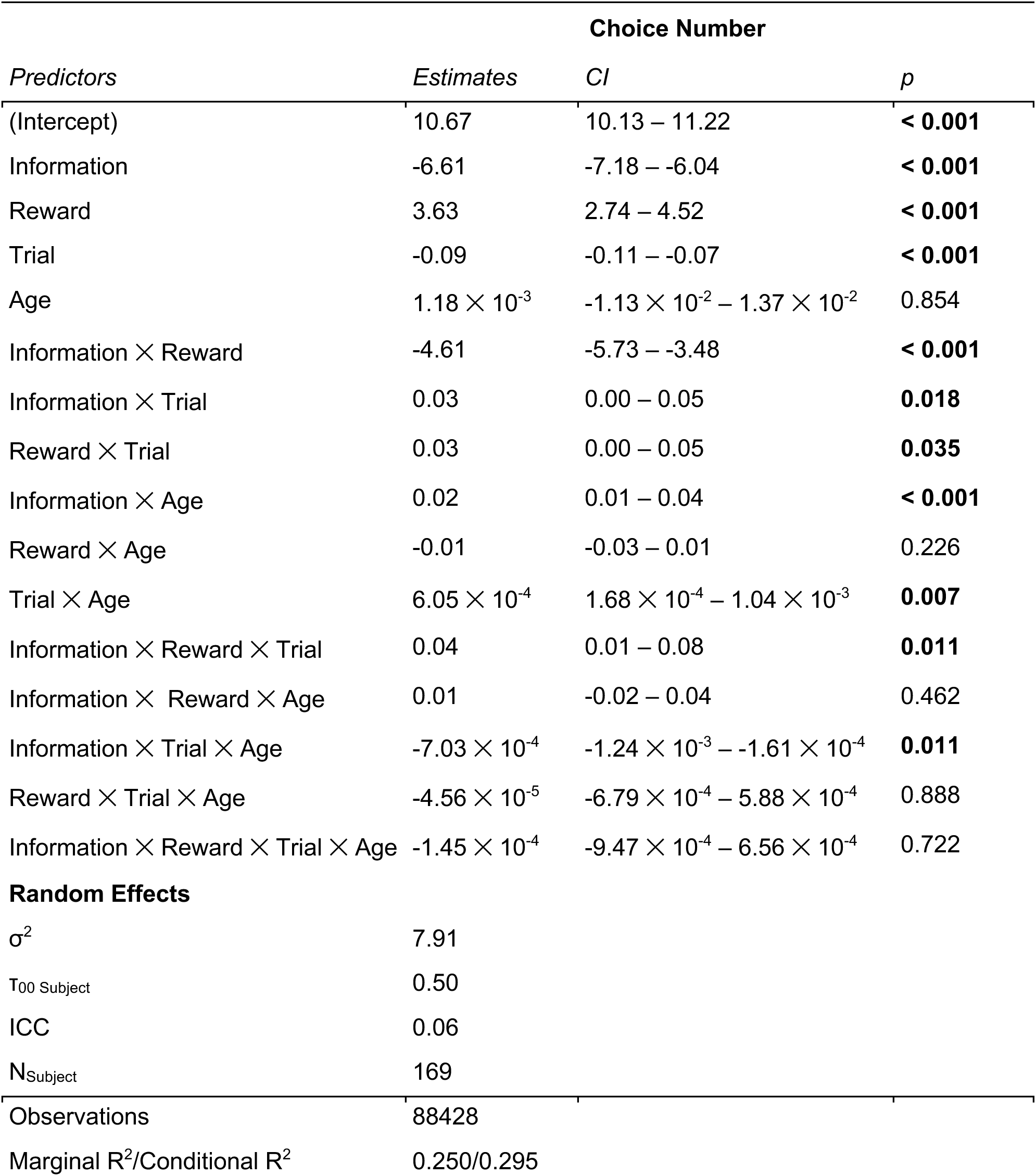
Information and Reward Mixed-Effects Model Results. The linear effects of information, reward, trial number, age, and all interactions on choice number within trial were examined.

Notably, unlike younger adults, older adults were motivated by information throughout the session. A significant three-way interaction between information, trial number, and age (see “(Information ✕ Trial) ✕ Age” in **Table 2**) was found. Adults prioritized information earlier in each trial, and this prioritization weakened as trial number increased. However, age affected this relationship. Whereas younger adults prioritized information less in later trials, older adults were driven by information throughout the duration of the task, even in late trials. This finding is evident in **Figure 3**: for younger participants, as trials progress from early trials (red line) to middle trials (green line) to late trials (blue line), the strength of the effect of information on choice number (slope) decreases. For older participants, however, the slope is relatively constant from early to late trials. Therefore, the three-way interaction between information, trial number, and age appears to be driven by the attenuated effect of information on choice in later trials in younger as compared to older adults.

### More exploration does not pay off

We next sought to understand what benefit heightened exploratory behavior may confer in older adults. In our battleship task, choice outcomes provide both information and reward, and as reported above, older adults are more driven by information, maintain their preference for information throughout the session, and engage in more directed exploration than younger adults. Thus, we sought signatures of motivated information seeking in participants’ choice behavior.

Previous findings using this task revealed foraging-type search strategies in both humans and non-human primates which related to behavior (Barack et al., 2023). Foraging is the capacity to search for and find resources under ignorance (Barack, 2024): an iterated series of decisions during search where the goal is at an unknown location. In foraging theory, decisions like where to search and when to switch locations are made by comparing current performance to the average across the environment (Charnov, 1976; Schmid-Hempel, 1988). To examine foraging behavior in the current study, we computed a custom forager score for each participant to determine search efficiency for information and reward (see Methods). Forager scores could not be calculated for 15 of 169 (∼9%) participants. A logistic regression model indicated no significant relationship between age and the likelihood of failing to calculate a forager score (OR = 1.02, 95% CI [0.987, 1.065], *p* = 0.267). Information forager score and reward forager score were significantly negatively correlated (r(154) =-0.629, *p* < 0.001). Moreover, correlations between age and each type of forager score revealed that increasing age was associated with higher information forager scores (r(154) = 0.282, *p* < 0.001) and lower reward forager scores (r(154) =-0.278, *p* < 0.001). These findings suggest that older adults forage more for information while younger adults forage more for reward.

However, a regression model assessing the effect of both forager scores and age on reward rate revealed that information forager score was negatively related to the reward rate (OLS main effect of info forager score β = −0.291, 95% CI [−0.458, −0.124], *p* < 0.001), while reward forager score was positively related to reward rate (OLS main effect of reward forager score β = 0.322, 95% CI [0.155, 0.488], *p* < 0.001). There were no interactions between either forager score and age on reward rate, indicating that the relationship between each forager score and reward rate did not change with age. This suggests that more efficient foraging for information in this task is detrimental to reward maximization across the session.

There may however be benefits to more efficient information foraging that are orthogonal to the receipt of reward. One possibility is that better foraging for information could help learn the shapes faster, even if reward rate remains low. We have previously posited that a changepoint detection algorithm applied to the cumulative choices over trials indexes shape learning (Barack et al., 2023) and demonstrated that an earlier changepoint was associated with a higher information foraging score. We largely replicate those findings here (see Supplementary Material). However, the changepoint detection method may instead index behavioral stabilization as described above rather than speed of learning, particularly in older adults. To newly assess speed of learning, we fit an exponential decay function to the number of choices made to reveal a shape across presentations of each shape. One of the parameters of the exponential function estimates the steepness of the decay in number of choices across presentations, a measure of the speed of learning a shape. Another parameter estimates the curve’s asymptote (i.e., the number of choices a participant continues to need once the shape has been fully learned), a measure of the stable level of performance reached for that shape. Bayesian mixed effects modeling then allowed us to estimate the interaction between the information foraging score and age on speed of learning and on projected final post-learning performance across shapes and individuals. The model revealed no credible interaction between info forager score and age on per-shape speed of learning (M = 0.026, SD = 0.081, 95% Highest Density Interval [HDI] = [−0.136, 0.179], BF_10_ = 0.17, which constitutes moderate evidence for the null), nor a credible main effect of age on speed of learning (M = −0.098, SD = 0.083, 95% HDI = [−0.264, 0.059], BF_10_ = 0.33, which constitutes moderate evidence for the null). The lack of a main effect of age on speed of learning suggests the observed slowdown was not due to age itself. There was however a credible negative main effect of information foraging score on speed of shape learning, slowing it down (M= −0.363, SD = 0.088, 95% HDI = [−0.53, −0.19], BF_10_ ≈ 900, which constitutes extreme evidence for an effect). There were also no credible main effects or interaction between age and info forager score on projected final stable per-shape performance (all BF_10_ ≤ 0.03, which all constitute strong evidence for the null). These results suggest that persistent foraging for information, regardless of age, slows the speed at which one learns shapes but does not predict the projected final performance after learning. We discuss below why older adults might persist in exploration and information seeking, even when it appears to decrease receipt of reward and speed of learning.

## Discussion

Older adults searched far more actively for information in our task than younger adults, despite achieving a lower overall point score. Older adults relied more heavily on the search for information, spreading their choices more widely, engaging in more directed exploration, making more choices per trial, and acquiring more information per trial than younger adults. Additionally, older adults showed a prolonged reliance on informative feedback over rewarding feedback when making new choices. Yet, this pronounced information seeking, regardless of age, resulted in slower learning and less reward. Why older adults persist in such effortful information-motivated search, at a cost of slower learning and a lower reward rate, is the central question we take up below.

Decades of research clearly show that older adults learn from reward feedback worse than younger individuals (Eppinger et al., 2013; Frank & Kong, 2008; Lerner et al., 2018; Mell et al., 2005). This learning deficit is largely due to changes in the dopaminergic system in the aging brain that is responsible for processing the experience of reward (Bäckman et al., 2000; Chowdhury et al., 2013; Eppinger et al., 2012, 2013; Kaasinen & Rinne, 2002; Samanez-Larkin et al., 2010, 2014; Schott et al., 2007; Volkow et al., 1998). Other work has identified the same dopaminergic system as key to processing so-called non-instrumental information — information that is not useful for guiding future behavior to obtain rewards (Bromberg-Martin & Hikosaka, 2009; Bromberg-Martin & Monosov, 2020; Sharot & Sunstein, 2020). These neural findings make sense of behavior that suggests human and non-human animals value and enjoy receiving information, just like they do reward (Blanchard et al., 2015; Bromberg-Martin et al., 2024, 2026; Bromberg-Martin & Hikosaka, 2009, 2011; Bussell et al., 2026; Charpentier et al., 2018). Put together, one might expect older adults to have deficits in learning from reward or information. Our findings only partly bear out this prediction: despite a pronounced preference for information, older adults did not learn the environment better or earn more reward. What is unexpected is not superior learning but the persistence of information seeking itself — why do older adults invest so heavily in gathering information when it yields no advantage?

In the battleship task, older adults appear to prioritize information over reward while searching for hidden shapes, even late in the experiment, presumably after they have learned the shapes and their locations. Younger adults, by contrast, appear to switch from exploratory information seeking to exploitative reward seeking. This finding runs counter to prior studies that suggest older adults exhibit less exploratory behavior when compared to their younger counterparts (Donnellan & Lucas, 2008; Spreng & Turner, 2021; Willig et al., 1987). But there are some notable exceptions. First, older adults outperform younger adults on a simple bandit task where the rewards are dependent on choice history (Worthy et al., 2011). Older adults accomplish this by adaptively deploying a win-stay lose-shift strategy, repeating choices (i.e., exploiting) when they win and switching (i.e., exploring) when they lose. Second, when given an opportunity to learn more facts about a current topic or learn about a new topic, older adults explore a topic more deeply, whereas younger adults explore more widely across topics (Fastrich et al., 2024). Finally, age is correlated with more exploration in callitrichid monkeys in zoos (Kendal et al., 2005). The current findings complement these observations of adaptive exploration in older adults and link greater visuospatial exploration to slower learning of features of a novel environment and a lower reward rate.

Greater exploration can be driven by a desire for information (i.e., directed exploration) or the product of random behavioral variability (i.e., random exploration, Wilson et al., 2021). While directed exploration has been shown to remain relatively stable across adulthood, random exploration has been shown to decrease (rather than increase) in older age (Mizell et al., 2024). We derived a measure of directed exploration from participants’ choices in our task by calculating the reward-and information-weighted distance from one choice to the next, computing the average weighted distance from one choice to all possible next choices, and then measuring the deviation from that average. This revealed that older adults were more directed in their exploration of the board, even in late choices of late trials, when compared to younger adults.

Although our analyses do not definitively establish an increase in directed exploration in older compared to younger adults, the pattern of exploration on our task provides additional evidence for this interpretation. Older adults were more likely to ‘jump’ (i.e., choose a more distant tile that is not connected to the last chosen tile) when the last tile yielded less information than the average across all choices, a pattern of search for instrumental information that conforms to the marginal value theorem in foraging theory (Charnov, 1976). Young adults, on the other hand, were more likely to jump when the last tile yielded less reward than the average. These results suggest, for the first time, that older adults prefer foraging for information compared to their younger counterparts. By contrast, younger adults prefer foraging for reward compared to older adults, as has been shown before (Wiegand & Wolfe, 2021). Why might adults prefer information foraging as they age when they were once keen foragers for reward?

Older adults show persistent directed exploration and greater information foraging throughout learning. The explanation for this pattern that we favor lies in the intrinsic, non-instrumental value of information, which increases with age: older adults seek information because information itself has become more valuable to them, not because it serves a downstream reward. How information is valued over the lifespan is little studied, and to our knowledge only one non-peer-reviewed study has shown older adults to be willing to incur a higher cost for information than young adults (Tedeschi, 2020). This suggests two non-exclusive mechanisms. 1) If individuals place increased value on information as they age, then their choices may be better driven by information than reward. The well-characterized deficiencies in dopaminergic processing (Bäckman et al., 2000; Chowdhury et al., 2013; Eppinger et al., 2012, 2013; Kaasinen & Rinne, 2002; Schott et al., 2007; Volkow et al., 1998) and the proposed role of dopamine in signaling non-instrumental informative outcomes (Bromberg-Martin & Hikosaka, 2009; Bromberg-Martin & Monosov, 2020) may seem to militate against this first option, since the same system that degrades with age also signals the value of information. This tension, however, need not be fatal to the account for two reasons: First, the information at stake in our task is instrumental — it guides choice toward reward — and the circuitry that values instrumental information may be comparatively spared in aging relative to the reward-processing system. Second, a decline in the value of reward would by itself raise the relative value of information; as reward valuation degrades with age, information comes to dominate choice even if its absolute value is unchanged. Our behavioral data cannot separate a rise in the value of information from a fall in the value of reward, but both predict the reallocation toward information that we observe. 2) Because informative options may be valued more highly in older adults, perhaps they consider the drop in information gain particularly aversive. Combined with the knowledge that older adults learn better from negative compared to positive feedback (Frank & Kong, 2008; Hämmerer et al., 2011), the comparative aversiveness of a drop in information from choices may drive older individuals to explore more (“jump” away from the last chosen tile), rendering them more efficient foragers for information. Older adults’ increased sensitivity to negative feedback in a learning context likely reflects lower dopamine levels, as shown in studies of clinical populations with altered dopamine (Frank et al., 2004; Palminteri et al., 2009). Higher information foraging in older adults would then be a byproduct of changes in the dopaminergic system of the aging brain — one whose consequences need not be adaptive, since the heightened pull toward information in our task impaired learning speed and lowered reward gains.

Overall, these findings offer a more nuanced take on learning in older adults than traditional reward-based research: rather than a simple deficit, we observe a reallocation of motivation toward information that persists despite measurable costs to learning and performance. That the drive to seek information persists despite the cost resonates with proposals that curiosity — the search for and prioritization of information — plays an increasingly central role in the aging mind (Sakaki et al., 2018). Why this pull toward information persists when it lowers reward and slows learning is, we suggest, the central puzzle these findings raise and understanding it may reshape how we engage and motivate aging learners.

## Materials and Methods

### Participants

Data were collected from 169 adults (*n* female = 86, *n* male = 81, *n* transgender = 2) ages 18-71 (*M* = 39; *SD* = 15.4) using a variety of recruitment and payment methods, namely SONA, an experiment management system, and CloudResearch’s MTurk Toolkit. Participants were required to speak fluent English, have no previous serious head injuries, have no neurological or psychiatric disorders, not be taking medications/drugs for psychiatric reasons, and have normal or corrected-to-normal vision and hearing abilities in order to participate in the study. Participants provided informed consent, and the study was conducted in accordance with the University of Chicago’s Institutional Review Board (IRB).

### Materials

The “Battleship” task created by Barack and colleagues (2023) was used (**Figure 1**). In this shape-search task, participants explored a 5×5 grid concealing one hidden shape made of multiple connected tiles. Their goal was to strategically select and “flip over” tiles to reveal the hidden shape using the fewest possible choices. Only one shape was hidden per trial. At the start of a trial, participants moved their mouse to a fixation target at the center of the screen (500 ms) until individual targets appeared over the remaining tiles. After a 500 ms delay, the central fixation square disappeared, allowing participants to move their mouse to a tile they wished to uncover. They held their position on the selected tile for 250 ms to reveal what was underneath. If a participant selected a tile that revealed part of the hidden shape, they received a reward (1 point) for this “hit.” If they selected an empty tile, or a “miss,” they were not rewarded. The participants’ point totals were visible to them throughout the duration of the task. The series of choices taken to uncover one shape constituted one complete trial. Once the shape was revealed, the board reset, a new shape was randomly chosen, and the participant began their search again.

This paradigm used five distinct shapes, each with a unique location at which it remained across all trials and participants (**Figure 1**). Shape color varied pseudorandomly from trial to trial. The shapes in this task were required to be made up of at least 3 tiles, but no more than 8 tiles. The tiles could not be only diagonally connected; to form a shape, tiles had to share at least one border with another tile. The number of possible fully revealed boards was 3904. This set of states (n_state_= 3904) was assumed to be known to participants in order to simplify entropy-related computations. This task was completed online for all participants.

Our task design allows information to be disambiguated from reward. Information is the difference in the entropy of the probability distribution over the set of shapes before and after each choice outcome. In this paradigm, each choice made by a participant reduces the possible state for that trial, thus reducing the entropy and providing information about which shape and where the reward might be. Rewards are the points delivered as outcome from selecting a tile (0 for a miss or 1 for a hit).

### Procedure

The present study was conducted entirely online. After being recruited through either the University of Chicago’s SONA system or Amazon Mechanical Turk (MTurk), participants were redirected through an online university server that hosted the task. Participants were immediately met with an online consent form detailing what the study would involve, compensation, and contact information. Upon providing consent via a mouse click, participants were taken to an instruction page detailing what the task was and how to complete it. A short (5-7 question) true or false quiz was used as an attention and comprehension check to ensure all participants understood how to complete the task before moving on. Once participants answered all questions correctly, they were taken to the task, where they repeated the process of tile selection and shape searching detailed above for a total of 50 minutes. At 50 minutes, the task stopped automatically, and participants completed a questionnaire, which asked questions about demographic characteristics (e.g., age, race) and how participants completed the task to better understand their strategies in learning the shape set. Participants were then given a completion code to either send to the researcher (in the case of SONA users) or input into MTurk, which the researcher would then verify. Once verified, participants were provided with compensation for completing the study. Participants in the young adult group were compensated with 1.5 course credits, and all other participants were compensated with $7.50.

### Determining time of behavioral stabilization

Behavioral stabilization during learning was operationalized using a changepoint detection test (Gallistel et al., 2001; Inclan & Tiao, 1994) on the mean and variance of choice. This test takes the cumulative sum of the number of choices used to find a shape and looks for changes in the rate of accumulation (Gallistel et al., 2001). The changepoint detection task was first used on the mean number of choices required to find a shape hidden on the grid. While participants are learning the shape set, they initially require a larger number of choices to uncover a shape, leading to a rapid rise in the cumulative sum. However, once participants have nearly learned or fully learned the shape set, the number of choices required to uncover a shape will be far less than at the start of the task, leading the cumulative sum to rise much slower. The cumulative sum is calculated at each trial, and the log-odds of a change in slope are computed and tested against a changepoint detection threshold (threshold = 4, corresponding to *p* < 0.001). The changepoint detection test then determines the last detected changepoint for the mean number of choices it takes to unveil a shape. The test was run on the set of trials for each unique shape separately.

The changepoint detection test was also applied to the variance in the choices made to uncover a shape. At the start of the task and during early learning, participants are expected to exhibit greater variance in the number of choices needed to find a shape, as many outcomes in early trials can be a result of chance alone. However, as participants learn the shape set, they will become less variable in the number of choices needed to finish a shape. After applying the changepoint detection test described above, the value of best-fit line through each detected changepoint interval was subtracted from the number of choices on each trial, resulting in a vector of residuals for the number of choices to finish revealing a shape. We then computed the variance over a moving window of 5 trials for these vectors and then ran the changepoint detection test on the cumulative sum of that variance (Inclan & Tiao, 1994).

The last changepoint (either by variance or by mean) for each participant marked the point at which their search behavior stabilized. This test failed to detect a changepoint for 13 out of 169 (∼8%) participants. These participants were excluded from changepoint analyses as a result.

### Quantifying choice distance

To quantify the spatial distribution of participants’ choices, we computed a choice distance metric based on pairwise Euclidean distances. For each participant, we calculated the Euclidean distances between all pairs of selected tiles in each trial. The mean choice distance for each trial was then derived, providing an average measure of how dispersed participants’ selections were. This allowed us to examine variations in dispersion across the task. Higher choice distance values indicate more widely distributed choices, while lower values suggest more clustered decision patterns.

### Quantifying directed exploration

To determine whether participants’ exploration reflected directed exploration rather than random exploration, we compared participants’ actual movements to an expected movement baseline. Directed exploration was operationalized as the deviation from the expected movement on each choice.

For each choice within a trial, we defined the expected movement as the average Euclidean distance from the previously selected tile to all remaining unchosen tiles for that trial. Because participants’ decisions are influenced not only by spatial factors but also by the expected reward and expected information of each tile, the Euclidean distances were weighted by both of these factors. For each choice, we computed two weighted averages: one in which all distances were multiplied by the expected reward and another in which distances were multiplied by the expected information. The mean of these two weighted averages was taken as the expected movement for that choice.

Selecting a tile near this average suggests that choices closely follow the expected movement, consistent with random exploratory behavior. In contrast, choosing a tile that deviates from this baseline indicates that participants select tiles in a manner that diverges from the simple expected pattern, reflecting a more goal-directed, systematic choice beyond random behavioral variability.

Directed exploration was therefore quantified as the absolute difference between this expected movement and the actual distance traveled by the participant on that choice.

### Quantifying information

Information was defined as the difference in the entropy before and after each choice outcome, with each choice reducing the possible number of configurations of the fully-revealed 5×5 grid. A distribution of probabilities for each revealed shape, the possible pattern of filled and empty tiles that the 5×5 grid could have once fully revealed, was used to compute information outcomes. In the following, ‘state’ is a possible configuration of filled and empty tiles on a fully revealed board. While there are 3,904 possible states of the board given the constraints on the shapes, there were only five states used for the task, one for each unique shape in the shape set.

We used a Dirichlet distribution to model the states across trials. To illustrate, imagine that each shape is allocated a “jar” that collects tokens across trials. Each time a specific shape is hidden during a trial (e.g., the purple square in **Figure 1**), the jar pertaining to the purple square receives a token. Each jar begins the task with two tokens and accumulates tokens across all trials as the state pertaining to each jar is revealed at the end of each trial. To ensure numerical stability and proper updating using the maximum *a posteriori* estimate of the Dirichlet distribution, two initial tokens (i.e., counts) are required for each state. Because the maximum *a posteriori* estimate subtracts one from each bin, and bins cannot be 0 or less, and in order to prevent a division by zero, 2 was chosen as the simplest, smallest, and model-free numerically stable initial count for each state. As trials progress, the token counts in each jar are updated: the initial counts for each state at the start of each additional trial are the updated counts from the end of the most recent trial. For example, if Jar 1 received 1 token at the end of Trial 1, at the beginning of Trial 2, Jar 1 would contain 3 tokens, while the other jars would still contain 2 tokens each. Therefore, each jar represents a vector of counts, reflecting how many times a participant has previously uncovered a particular shape throughout the trials.

In addition to being updated at the end of the trial to reflect which shape was revealed, the distribution of probabilities of the states is also updated throughout the trials. At the beginning of each trial, the distribution of probabilities of the states during trials, labeled ‘B’ (for belief) in these analyses, was set to the values in the Dirichlet distribution, the prior set of probabilities. After each tile selection during the trial, B was updated with a vector of 1’s and 0’s for the likelihoods of each state given the outcome of that choice. If a given state was consistent with the outcome (e.g., a purple square state and the tile selection of the top-left tile on the grid, **Figure 1**), it would be updated with a 1; otherwise, it would be updated with a 0. The probability of each state prior to that choice was then multiplied by the likelihood of the outcome to yield a new probability for the state. The resulting probabilities for each state were then used to attain a new B, the new prior for the next choice in the trial.

Information was then defined as the difference in the Shannon entropy, H, of B before and after each choice outcome:

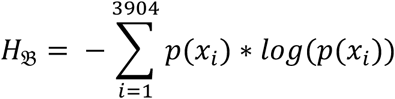

for the probability p(x_i_) of state x_i_. This definition was used in turn to define expected information:

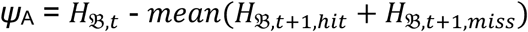

where ‘*ψ*_A_’ represents the expected information from selecting tile A and ‘*H*_*B*_’ is defined as above. *H*_*B*_ was computed for the current choice *t* and the mean *H*_*B*_ was calculated for the next choice *t*+1 assuming the choice of A resulted in a hit or a miss. Expected information was defined as the difference between the entropy before selecting A and the average entropy after selecting A.

This conceptualization of information reflects how much each hit or miss from a choice changes a participant’s uncertainty about the current trial’s state. In this way, even non-rewarding outcomes (i.e., misses) can yield information that can be used to move a participant closer to gaining reward and uncovering a shape.

### Quantifying reward

Reward was operationalized using a point system: for each “hit”, participants earned 1 point, contributing to an overall score visible throughout the task’s entirety. Expected reward for a given tile was defined by the number of times choosing that tile revealed a portion of the shape (hits) in the past divided by the total number of times that tile had been previously selected:

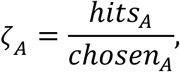

where ‘*ζ_A_*’ represents the expected reward of tile A, ‘hits_A_’ represents the number of times choosing A led to a reward, and ‘chosen_A_’ represents the number of times tile A had previously been chosen across all trials.

### Forager score

A custom foraging score was computed for each participant to assess their foraging abilities in terms of information and reward. Specifically, this score sought to gauge how efficient participants were at searching their environment for information and reward. Foraging decisions surround the serial presentation of decision options by which an animal must decide to exploit one option or move to search for a better option. This kind of decision-making differs from standard decision-making paradigms utilized in the literature, paradigms that revolve around binary, simultaneously-presented options. Moreover, in these foraging decisions, once-rejected options can be later returned to by foragers. A classic example of foraging behavior is demonstrated in patch leaving paradigms. In a new environment, animals must explore and make decisions about where to spend energy searching for resources in a way that is the most energetically favorable. An animal is expected to stay in a given area, or patch, until the current reward from that area has dropped below a threshold given by the opportunity cost of time, the opportunity cost of time being the long-run average reward per timestep (the marginal value theorem [MVT], Charnov, 1976; Constantino & Daw, 2015). In other words, taking the time to stay at a location and “harvest” involves missing out on possibly extremely rich other patches. Once the current, immediate reward is less than the accumulated average reward of the surrounding environment, the animal should choose to leave and search for a new patch (Charnov, 1976; Constantino & Daw, 2015). In the present study, a resource, or patch, was defined as a connected set of tiles. A “jump” indicated the choice to leave a given resource, or patch, and was defined as the choice of a non-neighboring tile, the choice of a tile more than one tile away from the most recently-chosen tile.

To calculate these scores, we examined each sequence of three neighboring tile choices followed by a jump, and we considered three features of foraging behavior. First, while participants decide to stay at a certain region of the board, staying within a given patch, the information or reward gained from the prior choices should be above the environmental average. Second, the choice outcomes preceding a “jump” decision, or the decision to leave a given resource, should be below the environmental reward or information average. Third, the choice outcomes preceding stay decisions should be greater than those preceding leave decisions. We computed two different types of forager scores, one for information foraging abilities and one for reward foraging abilities. **Figure 4** depicts this scoring process.

**Figure 4.**
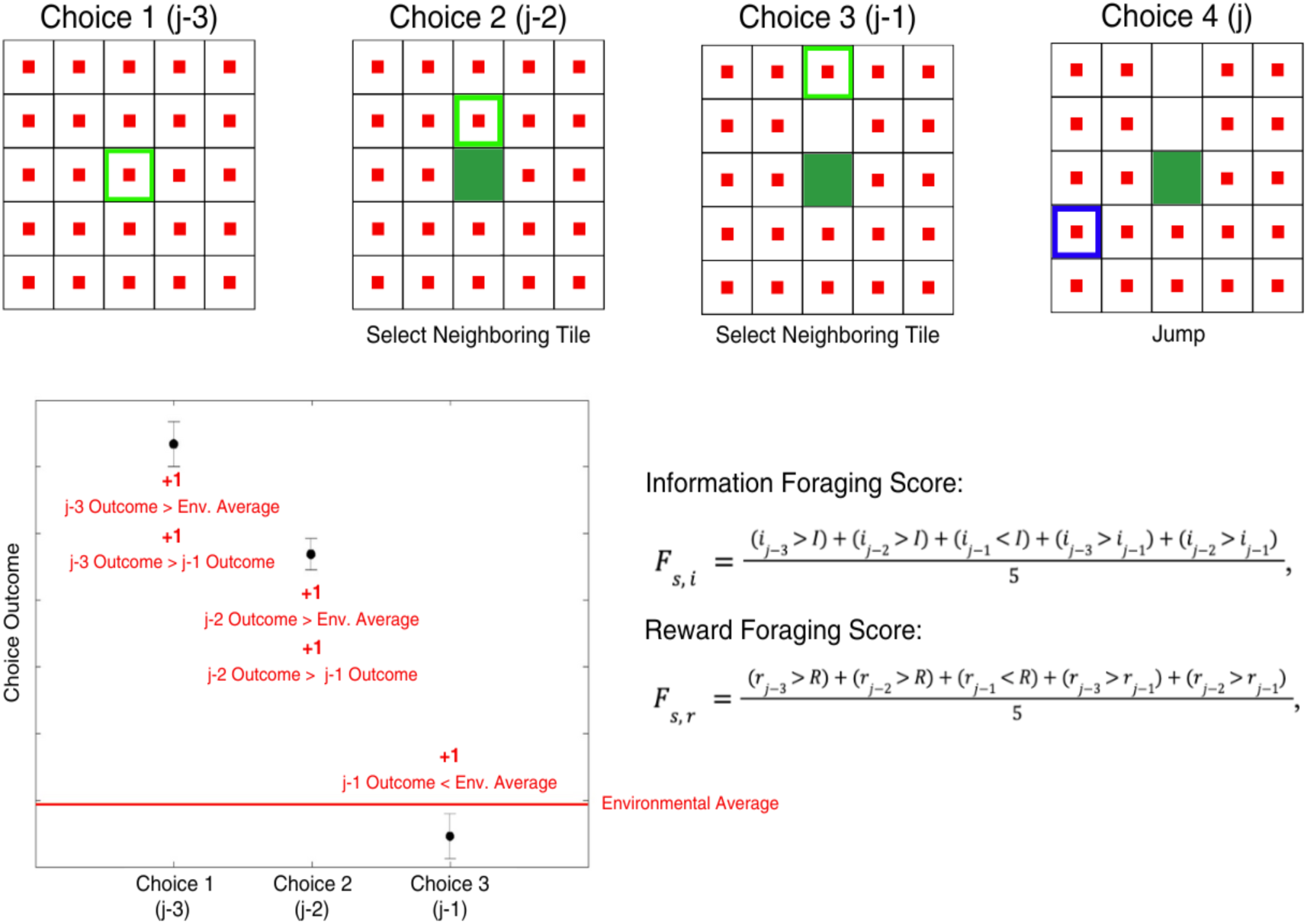
Calculating Forager Scores. An example of choice patterns examined by the forager is shown. Choice patterns with three consecutive neighboring tile choices followed by the selection of a non-neighboring tile (i.e., “jump”) were examined. Forager scores for information and reward were calculated separately for each participant. Forager scores were not found for 15 adults; these participants were excluded from all analyses involving these scores.

We calculated the information forager score in the following way:

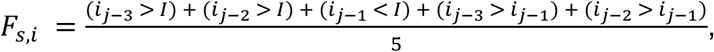

where *F_s,i_* is the information forager score for a given subject *s*, *i_j-3_* is the information outcome 3 choices prior to the jump, *i_j-2_* is the information outcome 2 choices prior to the jump, *i_j-1_* is the information outcome 1 choice prior to the jump, and *I* is the average information outcome in the environment. The reward forager score was calculated in a similar manner:

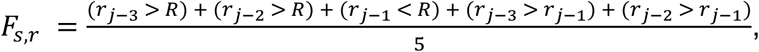

where *F_s,r_* is the reward forager score for a given subject *s*, *r_j-3_* is the reward outcome 3 choices prior to the jump, *r_j-2_* is the reward outcome 2 choices prior to the jump, *r_j-1_* is the reward outcome 1 choice prior to the jump, and *R* is the average reward outcome in the environment.

*I* and *R* were calculated separately for each participant by taking the average of all of the information or reward outcomes during learning. A score of 1 was the maximum forager score possible, as in order to be a perfect forager in the sequences of choices examined, the two choice outcomes prior to the jump must be above the environmental average (+2), the choice outcome directly preceding a jump must be below the average (+1), and the two choice outcomes prior to the pre-jump outcome must be greater than the pre-jump outcome (+2). See **Figure 4**. Each forager score was computed for every sequence of three choices of a neighboring tile prior to a jump; information foraging scores and reward foraging scores were then averaged by participant. We then entered these scores into regressions.

Forager scores were not found for 15 of 169 (∼9%) adults. This is due to these participants having no trials with the choice pattern utilized in calculating the forager score (i.e., 3 consecutive neighboring tile choices followed by the selection of a non-neighboring tile). These were the same adults by which the changepoint detection test failed in addition to two more participants. Those adults were excluded from all analyses involving these scores.

### Primary analyses

To explore how adults of varying ages completed the task differently in general, we employed ordinary least squares (OLS) linear regressions to examine the relationships between (1) total points earned and age, (2) final changepoint and age, (3) total number of trials completed and age, (4) total number of choices made and age, (5) average number of choices per trial and age, and (6) average information per trial and age. To account for multiple comparisons, all reported results were corrected using the false discovery rate (FDR) procedure.

To test whether the changepoint detection was capturing a genuine shift in learning rather than a point where older adults became satisfied with not knowing, we asked whether age influenced changes in choice variance before and after the changepoint. Choice variance was computed across a moving window of five trials. We then fit a linear mixed-effects model with variance as the outcome and the hidden shape, mean shape size across the moving window, learning phase (pre-or post-changepoint), age, and the interaction between age and learning phase as predictors. Random intercepts accounted for individual participant differences.

As a changepoint was not detected for 13 of the 169 participants, we ran a logistic regression to test whether age predicted successful changepoint detection. The dependent variable was binary, indicating whether a participant’s changepoint was successfully detected, and age was included as a continuous predictor.

We also sought to better understand how age influenced the spread, or distribution, of participants’ choices. Thus, we performed a mixed-effects linear regression to regress the Euclidean pairwise distance between choices against age, choice number in trial, and trial number in session, as well as all possible interactions (see **Table 1**). The model accounted for individual variability by including random intercepts for participants.

To examine how age influenced directed exploration, as opposed to random exploration, we regressed both measures (see above) against choice number, trial number, age, and all interactions against each measure in a mixed-effects linear regression that included random intercepts per participant.

To investigate how adults across the age span utilize information and reward to guide choices and learning, we performed a mixed-effects linear regression to regress choice number in trial against trial number in session, the expected reward for the chosen tile, the expected information for the chosen tile, and age. This model also tested for all possible interactions between trial number in session, expected reward for the chosen tile, expected information for the chosen tile, and age (see **Table 2**). For random effects, we included intercepts for subjects due to an insufficient number of trials to estimate random slopes. Mixed-effects linear regressions were also employed to examine the relationships between (1) information and reward and (2) information and reward across trials.

We used the “lme4” R software package (Bates et al., 2015) to perform mixed effects linear regressions along with “lmertest” (Kuznetsova et al., 2017), which used the Satterthwaite method (Satterthwaite, 1946), for estimating the p-values and degrees of freedom for these models.

We next considered how adults of varying ages foraged their environments to learn the shape set. A Pearson correlation test was run to examine the relationship between information forager score and reward forager score. Spearman correlation was employed to examine the relationships between (1) information forager score and age, (2) reward forager score and age.

As a forager score was not detected for 15 of the 169 participants, we ran a logistic regression to test whether age affected the calculation of a forager score. The dependent variable was binary, indicating whether a participant’s forager score was successfully calculated, and age was included as a continuous predictor.

To characterize how quickly and accurately each shape was learned, we modeled the number of choices required to reveal a shape as a function of the number of times that shape had been presented. For each participant and each of the five shapes, we ordered the trials on which that shape appeared and indexed them by presentation number. Only (participant, shape) series with at least five presentations were included, and analyses were restricted to the 154 participants for whom forager scores could be computed. The number of choices on presentation x was modeled with an exponential decay function:

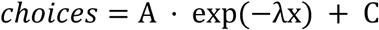

whose parameters decompose learning into conceptually distinct components. The rate λ captures the steepness of the decline in choices across presentations (i.e., the speed of learning a shape), with larger values indicating faster learning. The asymptote C is the value the curve is projected to settle toward as presentations continue. Because it extrapolates beyond the presentations actually observed, it indexes the projected final level of performance a participant would reach once the shape is fully learned, rather than an endpoint necessarily attained within the fixed-time session. The amplitude A captures the total reducible improvement from the first presentation to that projected asymptote, such that A + C is the expected number of choices on the first presentation of a shape.

We estimated these parameters within a single Bayesian nonlinear mixed-effects model fit jointly to all observations, rather than fitting each curve separately, so that information is pooled across participants and shapes and uncertainty in the curve parameters is propagated into all downstream inferences. Because each parameter is strictly positive, each was modeled on the log scale. For every parameter, the log-scale value for a given observation was the sum of a population-level intercept, fixed effects of standardized age, standardized information forager score, and their interaction, a reference-coded fixed effect of shape identity (five levels), and a participant-specific random effect; residual choices were modeled as Gaussian with a single standard deviation. The same model structure was used to estimate the effect of age alone on each parameter and the age-by-information-forager-score interaction reported in the main text. Random effects used a non-centered parameterization. We placed weakly informative priors throughout: normal priors on the log-scale intercepts centered on plausible values (rate, log 0.2; amplitude, log 4; asymptote, log 5), Normal(0, 0.5) priors on all age, forager-score, interaction, and shape coefficients, and half-normal priors on the random-effect and residual standard deviations. Age and information forager score were z-scored, so all corresponding coefficients are expressed per standard deviation.

Posterior distributions were sampled with the No-U-Turn Sampler (NUTS) implemented in NumPyro (Phan et al., 2019), using four chains of 1,000 warmup and 1,000 post-warmup iterations each (4,000 retained draws) and a target acceptance probability of 0.9. Convergence was confirmed by R^ ≤ 1.01 and adequate effective sample sizes for all population-level parameters, with no divergent transitions. For each population-level effect we report the posterior mean, the posterior standard deviation, and the 95% highest-density interval (HDI). To quantify evidence for and against each effect, we computed Savage–Dickey density-ratio Bayes factors (BF₁₀), comparing the posterior density at zero to the prior density at zero under the Normal(0, 0.5) prior; Bayes factors were interpreted using conventional thresholds, with values (or their reciprocals, BF₀₁) between 3 and 10 taken as moderate evidence, 10–30 as strong, 30–100 as very strong, and greater than 100 as extreme.

## Author Contributions

A.H.: conceptualization, methodology, software, investigation, data curation, formal analysis, writing - original draft, writing - review & editing, visualization, project administration; D.L.B.: conceptualization, methodology, software, writing - original draft, writing - review & editing; A.B.: resources, conceptualization, methodology, software, validation, formal analysis, writing - original draft, writing - review & editing, supervision, funding acquisition.

## Acknowledgements

This work was made possible by the support of Grant 2342775 from the U.S. National Science Foundation (A.B.), and Grant 63750 from the John Templeton Foundation (D.L.B.). The opinions expressed in this publication are those of the researchers and do not necessarily reflect the views of the funders.

