## Supplemental Material for "Older Adults Forage Broadly for Information Despite Cost"

### Supplemental results

*Better search for information leads to earlier behavioral stabilization*

The ability to forage for information was associated with an earlier changepoint (i.e., earlier behavioral stabilization; OLS β = -63.149, 95% CI [-107.744, -18.553], $p$ = 0.006). However, while age and information forager score were positively correlated, there was no main effect of age or interaction between age and information forager score on changepoint (**Figure S1**). In other words, better ability to forage for information was associated with earlier behavioral stabilization, but this effect did not differ significantly based on age, even though age predicted information forager scores. In contrast, participants who were more efficient reward foragers stabilized their search behavior later than those who were less efficient, as the ability to forage for reward was associated with later final changepoints (β = 127.684, 95% CI [16.278, 239.089], $p$ = 0.025). This effect of reward foraging on final changepoint also did not differ significantly based on age (**Figure S1**).

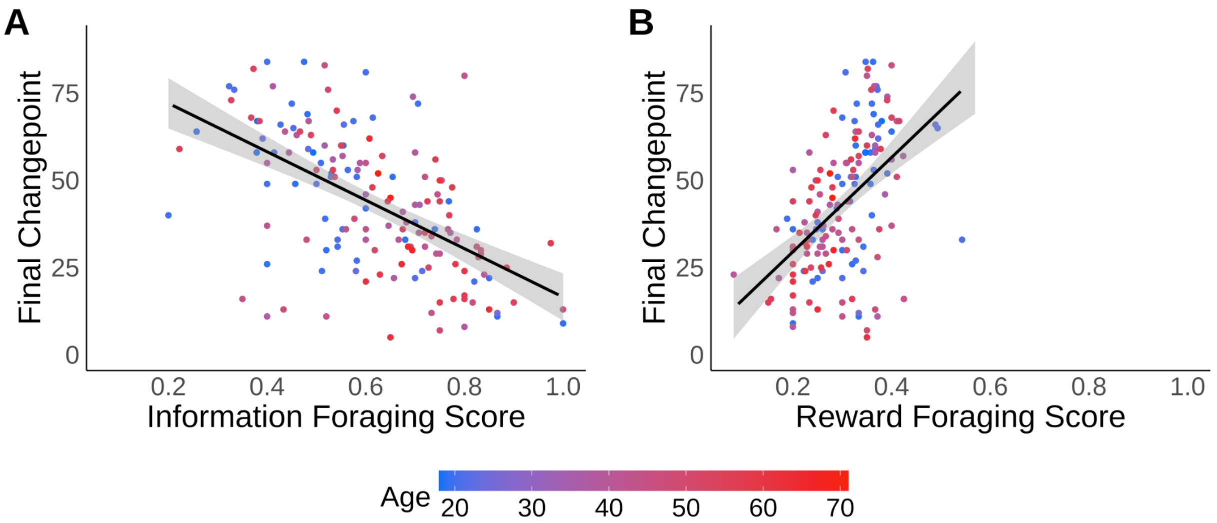


**Figure S1. Timing of Behavioral Stabilization and Forager Scores are Correlated.** Last changepoint was regressed against **A)** information forager score and **B)** reward forager scores. Raw data points for each participants' forager score and changepoint are plotted. Participant age is indicated by color (youngest in blue to oldest in red). Lines are fits of linear regression. Shaded areas are 95% confidence bands.
